# Differential receptor crosstalk in DRD1-DRD2 heterodimer upon phasic and tonic dopamine signals

**DOI:** 10.1101/2021.05.13.444104

**Authors:** Hyunbin Kim, Min-Ho Nam, Sohyeon Jeong, Hyowon Lee, Soo-Jin Oh, Jeongjin Kim, Nakwon Choi, Jihye Seong

**Author notes:** To whom correspondence should be addressed., Jihye Seong, Ph.D., Brain Science Institute, Korea Institute of Science and Technology (KIST), Seoul, Republic of Korea.

## Abstract

In response to phasic and tonic release, dopamine neurotransmission is regulated by its receptor subtypes, mainly dopamine receptor type 1 and 2 (DRD1 and DRD2). These dopamine receptors are known to form a heterodimer, however the receptor crosstalk between DRD1 and DRD2 was only suspected by measuring their downstream signaling products, due to the lack of methodology for selectively detecting individual activity of different dopamine receptors. Here, we develop red DRD1 sensor (R-DRD1) and green DRD2 sensor (G-DRD2) which can specifically monitor the real-time activity of DRD1 and DRD2, and apply these multicolor sensors to directly measure the receptor crosstalk in the DRD1-DRD2 heterodimer. Surprisingly, we discover that DRD1 activation in the heterodimer is inhibited only at micromolar phasic concentration of dopamine, while DRD2 activation is selectively inhibited at nanomolar tonic dopamine level. Differential receptor crosstalk in the DRD1-DRD2 heterodimer further modulates their downstream cAMP level. These data imply a novel function of the DRD1-DRD2 heterodimer at physiological dopamine levels of phasic and tonic release. Our approach utilizing multicolor receptor sensors will be useful to discover novel function of GPCR heterodimers.

## Introduction

Dopamine is an important neurotransmitter which regulates motivation, locomotion and reward process (*1, 2*). Dysregulation of dopaminergic pathway thus causes neurological diseases such as Parkinson’s disease, schizophrenia and depression (*3, 4*). Dopamine binds to its receptors and the activated dopamine receptors mediate dopaminergic signaling pathways. Dopamine receptors are a type of G protein-coupled receptor (GPCR), which is activated by binding to its extracellular ligands and initiates various intracellular signaling pathways by recruiting different G alpha proteins such as Gs, Gi, Gq, and G12/13 (*5*). Five subtypes of dopamine receptors have been discovered, i.e. DRD1, DRD2, DRD3, DRD4 and DRD5, and they couple with different G proteins and induce distinct downstream signaling events (*2, 6*). Complex dopaminergic pathways are modulated by different subtypes of dopamine receptors, thus it is crucial to understand the distinct functions of dopamine receptor subtypes.

DRD1 and DRD2 are two major subtypes of dopamine receptors (*7*). DRD1 shows wide expression patterns throughout brain, including striatum, nucleus accumbens (NAc), substantia nigra (SN), amygdala and frontal cortex, and also at a lower level in the hippocampus, thalamus and cerebellum (*4, 8*). DRD2 is expressed in striatum, SN, ventral tegmental area (VTA), hippocampus, cortical area and hypothalamus (*4, 8*). DRD1 is activated upon binding of dopamine and recruits Gs protein, which then activates adenylyl cyclase (AC) (*9*). The activated AC generates cAMP for next downstream events such as protein kinase A (PKA) activation and cAMP-mediated gene expression (*10*). In contrast, upon the binding of dopamine, activated DRD2 interacts with Gi which inhibits AC and reduces cellular cAMP levels (*9*). Thus, the activation of DRD1 and DRD2 can induce opposite effects on cAMP production. In addition, released G beta-gamma proteins subsequent to the Gi activation activate G protein-coupled inwardly-rectifying potassium channels (GIRK) (*11*), inducing the hyperpolarization of the neuronal membrane potential to inhibit the neuronal activity, which is also a distinct effect from Gs.

As activated DRD1 and DRD2 have opposite effects on cAMP production (*12*), they may control the activity of each other to balance the dopamine-mediated cellular functions. Indeed, some brain regions such as striatum and SN display overlapping expressions of DRD1 and DRD2 (*13–15*), and interestingly, DRD1 and DRD2 are known to form a heterodimer (*16–20*). It has been reported that the DRD1-DRD2 heterodimer can induce the signaling related to Gq, for example calmodulin-dependent protein kinase II (CaMKII) and brain-derived neurotrophic factor (BDNF) expression (*21*). The DRD1-DRD2 heterodimer and this modulated downstream signaling thus may contribute to the neuroplasticity changes in dopaminergic circuits (*19*).

Emerging evidence suggest the existence of various GPCR heterodimers and their novel molecular and cellular functions (*22–24*). Indeed, in addition to the DRD1-DRD2 heterodimer, it has been suggested that DRD1 can crosstalk with other GPCRs such as DRD3 (*25*) and adenosine receptor 1 (*26*); and DRD2 can interact with DRD3 (*27*), adenosine receptor 2 (A2A) (*28*) and 5HT2A (*29*). Therefore, it is important to investigate the novel functions or receptor crosstalk of the heterodimers as well as individual GPCR activities. Various methods have been applied to demonstrate physical interactions in these GPCR heterodimers, for example fluorescence resonance energy transfer (FRET), bioluminescence resonance energy transfer (BRET) and co-immunoprecipitation (co-IP) (*14, 30, 31*). For studying the functional crosstalk within the GPCR heterodimers, the downstream G protein signaling events have been measured by luciferase-based cAMP assay or fluorescent calcium assay (*16*). As such, receptor functional crosstalk with each other within GPCR heterodimers was not directly measured but only suspected by detecting these downstream signaling products.

Recently, green dopamine sensors have been developed by inserting a circular permuted fluorescent protein (cpFP) in the third intracellular loop (ICL3) of dopamine receptors, which can increase the green fluorescence in the active conformation of dopamine receptors upon dopamine binding (*32, 33*). The same groups also reported red dopamine receptor sensors (*34, 35*). However, these dopamine sensors include the multiple modifications even in the receptor part to maximize the dopamine binding affinity and sensor performance, which are useful for the sensitive detection of dopamine levels in vivo, rather than representing specific individual receptor activity of DRD1 and DRD2. Therefore, in this study, we develop novel red DRD1 sensor (R-DRD1) and green DRD2 sensor (G-DRD2), which specifically monitor individual activities of DRD1 and DRD2 keeping their natural dopamine binding affinity.

We apply these multicolor DRD1 and DRD2 sensors to directly measure the receptor crosstalk in DRD1-DRD2 heterodimer for the first time. Utilizing the R-DRD1 and G-DRD2, we surprisingly discover differential receptor crosstalk in the DRD1-DRD2 heterodimer depending on dopamine concentrations: DRD1 activation in the heterodimer is inhibited only at micromolar dopamine levels, while DRD2 activation is selectively inhibited at nanomolar dopamine concentration. Differential receptor crosstalk in the DRD1-DRD2 heterodimer further modulate their downstream cAMP levels. As tonic and phasic dopamine releases are in the range of nanomolar and micromolar levels, these data imply a novel function of the DRD1-DRD2 heterodimer at physiological dopamine levels of phasic and tonic release. These multicolor DRD1 and DRD2 sensors can be further applied to investigate complex dopaminergic signaling pathways in the brain.

## Results

### Development of red DRD1 sensor

To develop a red fluorescent DRD1 sensor, we first obtained circular permutated mApple (cpmApple) and the linker sequence (LSSPV-cpmApple-TRDDL) from jRGECO1a, an intensity-based genetically encoded calcium sensor (*36*). These sequences were inserted into the third intracellular loop of DRD1 to generate the prototype DRD1 red sensor (Fig. 1A). In this design, the binding of dopamine to DRD1 will induce an increase of red fluorescence intensity, thus we can monitor real-time DRD1 activation (Fig. 1B).

**Figure 1.**
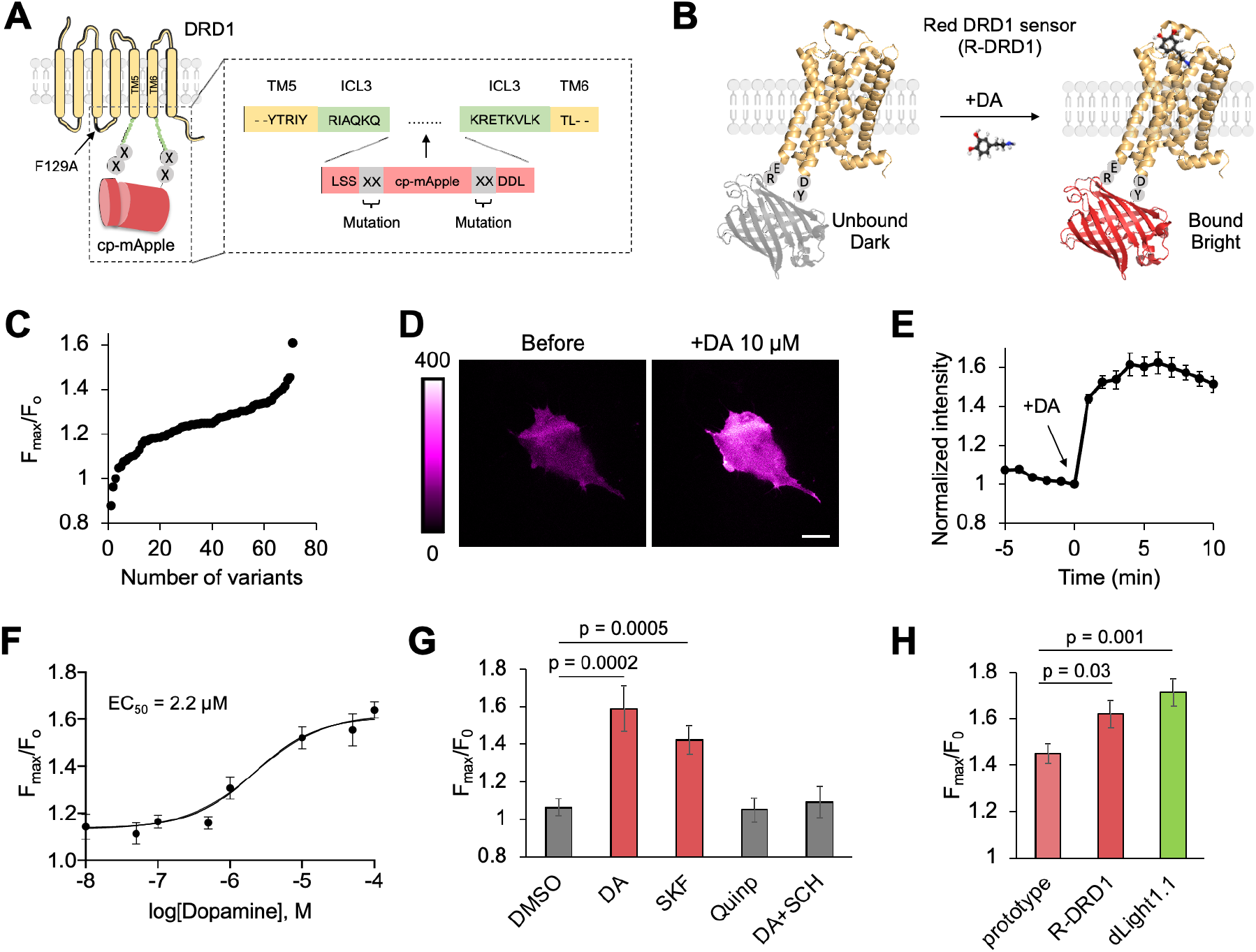
Development and characterization of R-DRD1. (**A**) Development strategy for R-DRD1. The prototype of R-DRD1 is generated by insertion of cpmApple with linker sequences (LSSPV-cpmApple-TRDDL) to the intracellular loop 3 (ICL3) of DRD1. Two amino acids in the N- and C-terminal of cpmApple were applied for the site saturation mutagenesis (shown as X). (**B**) Schematic of the R-DRD1 before and after the treatment of dopamine (DA). Red fluorescence of cpmApple in the R-DRD1 increases upon dopamine binding. (**C**) Screening for linker optimization. In HEK293A cells expressing different variants, normalized maximal intensity (*F*_max_ /*F*_0_) was compared after the treatment of 10 μM dopamine. Each dot represents average intensity change of each variant from more than 3 cells. (**D**) Representative images of the R-DRD1 before and after the treatment of 10 μM dopamine. The magenta color bar on left represents the range of fluorescent intensity. Scale bar, 20 μm. (**E**) Time course of normalized fluorescent intensity of R-DRD1 upon the treatment of 10 μM dopamine (n = 10). Data represent the mean value ± s.e.m. (**F**) *F*_max_ /*F*_0_ of R-DRD1 in response to different dopamine concentrations. EC_50_ was calculated as 2.2 μM. (**G**) Specificity of R-DRD1. *F*_max_ /*F*_0_ in response to DMSO (n = 11), dopamine (10 μM, n = 7), a DRD1 specific agonist SKF81297 (10 μM, n = 10), a DRD2 specific agonist quinpirole (10 μM, n = 4), and dopamine and a DRD1 antagonist SCH23390 (10 μM each, n = 5). Data are shown as means ± s.e.m. and p values were calculated by unpaired Student’s t-test. (**H**) *F*_max_ /*F*_0_ of the prototype (n = 13), R-DRD1 (n = 10), and dLight1.1 (n = 14) (*32*). Data are shown as means ± s.e.m. and p values were calculated by unpaired Student’s t-test.

Based on the prototype DRD1 red sensor, we also performed saturated random mutagenesis on the linker sequence, and the fluorescent intensity changes of these candidates after the treatment with 10 μM dopamine were screened in HEK293A cells (Fig. 1C). We found a candidate for a DRD1 red sensor which has a dopamine-induced increase of its red intensity (*F*_max_ /*F*_0_ = 145 ± 5%), and the selected sequences were LSSER-cpmApple-YDDDL inserted in the ICL3 of DRD1. We additionally introduced a mutation of F129A, which is crucial for interaction with G protein (*37*), and the dopamine-induced intensity change of the construct was further improved (*F*_max_ /*F*_0_ = 162 ± 6%) (Fig. 1C). This construct, LSSER-cpmApple-YDDDL with F129A, also shows an improved speed of the dopamine-induced response (fig. S1A). We named it as red DRD1 sensor or R-DRD1 (Fig. 1B, fig. S2).

R-DRD1 is well located at the plasma membrane in HEK 293A cells, and the real-time increase of red fluorescence intensity after the treatment with 10 μM dopamine can be observed by live-cell imaging under fluorescent microscopy (Fig. 1D-E, Movie S1). When we tested the response of R-DRD1 to different dopamine concentrations, the EC_50_ of R-DRD1 activation was 2.2 μM (Fig. 1F). R-DRD1 specifically responded to dopamine or a DRD1 specific agonist SKF81297 (SKF), but not by a DRD2 specific agonist quinpirole (Quinp) (Fig. 1G). In addition, the dopamine-induced response of R-DRD1 was prevented by a DRD1 specific inhibitor SCH23390 (SCH) (Fig. 1G). Additionally, we tested the possibility of triggering downstream signaling by R-DRD1 sensor expression at HEK293A cell. Using Glosensor (*38*), cAMP measuring assay which based on a luminescence, Wild type DRD1 transfected cells show significant cAMP response upon dopamine treatment, whereas R-DRD1 transfected cells show no significant cAMP response (fig. S1B). These results validate that R-DRD1 specifically reports the activity of DRD1 without interfering endogenous G protein signaling.

The dopamine-induced red intensity change of R-DRD1 was comparable to dLight1.1 (*32*) (*F*_max_ /*F*_0_ = 171 ± 6%), which is a previously reported green dopamine sensor based on DRD1 (Fig. 1H). The same group recently reported a red dopamine sensor called RdLight1(*34*). When we compared RdLight1 with R-DRD1, RdLight1 had a slightly larger intensity change compared to R-DRD1 (fig. S1C), but it also showed a considerable response to quinpirole, a DRD2 specific agonist (fig. S1D). In addition, the EC_50_ of RdLight1 is 859 nM while the one of R-DRD1 is 2.2 μM, suggesting the RdLight1 is a very sensitive dopamine sensor but the dopamine binding affinity for R-DRD1 is more similar to DRD1 (*39, 40*). Thus, R-DRD1 is a new red sensor which can specifically report the real-time DRD1 activity with its natural binding affinity.

### Development of green DRD2 sensor

To develop green DRD2 sensor, we first generated the prototype of DRD2 green sensor by inserting the linker and cpGFP module (LSSLI-cpGFP-NHDQL) from dLight1.1 (*32*) into the ICL3 of DRD2 (Fig. 2A). In this design, the binding of dopamine to DRD2 causes an increase of green fluorescence intensity, allowing the real-time visualization of DRD2 activation in live cells (Fig. 2B). The prototype DRD2 green sensor showed improved response (*F*_max_ /*F*_0_ = 209 ± 11 %) compared to GRAB_DA1m_, a previously reported DRD2-based green sensor (*33*) (Fig. 2C). We further improved the linker sequence by deleting one of the repeated leucine in the N-terminal side of cpGFP, which showed an approximately 50% increased response (*F*_max_ /*F*_0_ = 240 ± 10 %) comparing to the prototype green sensor (Fig. 2C). We named this version as green DRD2 sensor or G-DRD2 (Fig. 2B, fig. S2).

**Figure 2.**
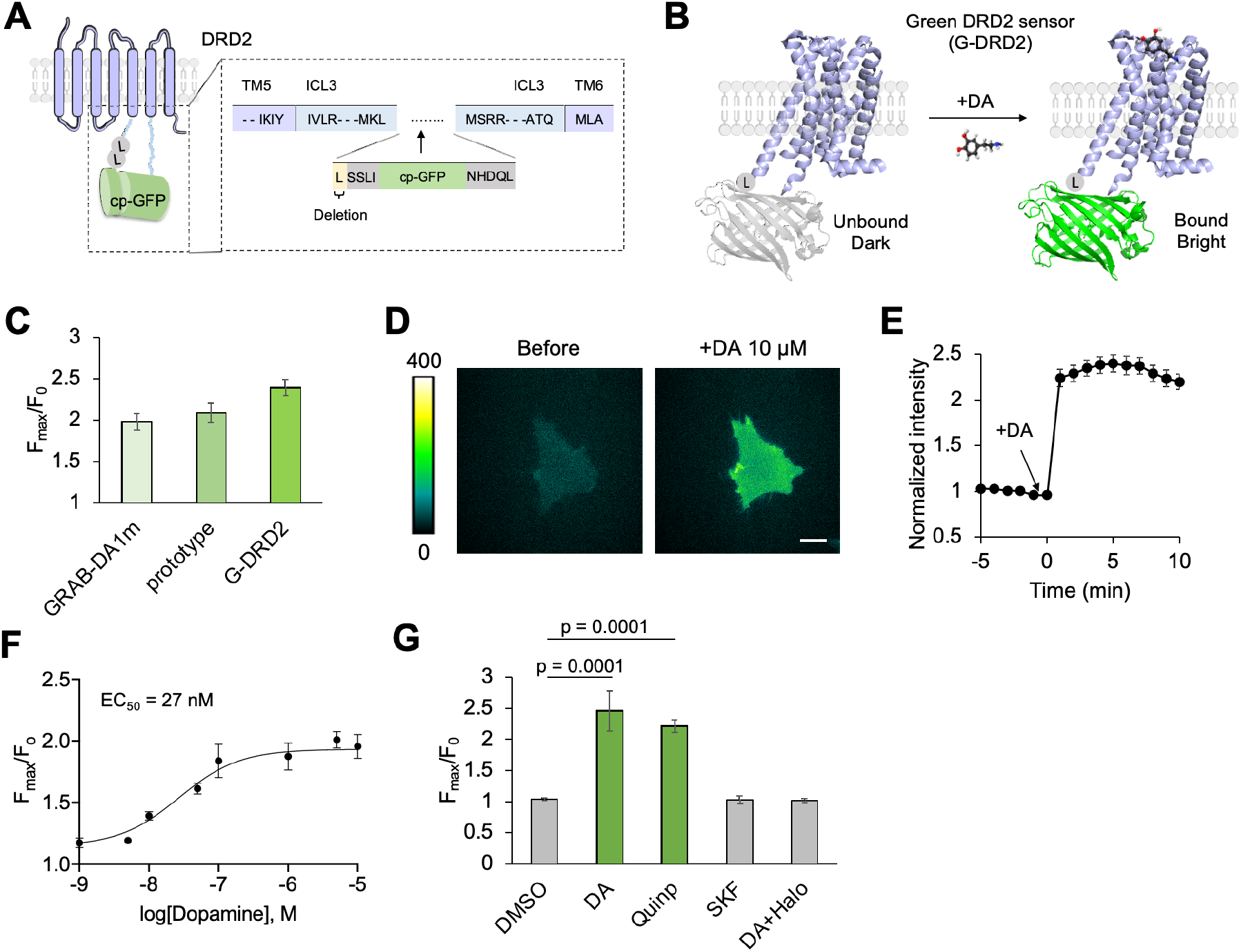
Development and characterization of G-DRD2. (**A**) Development strategy for G-DRD2. The prototype of G-DRD2 is generated by insertion of cpGFP with linker sequences (SSLI-cpGFP-NHDDL) to the intracellular loop 3 (ICL3) of DRD2. (**B**) Schematic of the G-DRD2 before and after the treatment of dopamine (DA). Green fluorescence of cpGFP in the G-DRD2 increases upon dopamine binding. (**C**) *F*_max_ /*F*_0_ of the GRAB-DA1m (n = 9) (*33*), prototype (n = 18), and G-DRD2 (n = 21). Data are shown as means ± s.e.m. (**D**) Representative images of the G-DRD2 before and after the treatment of 10 μM dopamine. The green color bar on left represents the range of fluorescent intensity. Scale bar, 20 μm. (**E**) Time course of normalized fluorescent intensity of G-DRD2 upon the treatment of 10 μM dopamine (n = 21). Data represent the mean value ± s.e.m. (**F**) *F*_max_ /*F*_0_ of G-DRD2 in response to different dopamine concentrations. EC_50_ was calculated as 27 nM. (**G**) Specificity of R-DRD1. *F*_max_ /*F*_0_ in response to DMSO (n = 11), dopamine (10 μM, n = 7), the DRD2 specific agonist quinpirole (10 μM, n = 5), the DRD1 specific agonist SKF81297 (10 μM, n = 4), and dopamine with a DRD2 specific antagonist haloperidol (10 μM each, n = 5). Data are shown as means ± s.e.m. and p values were calculated by unpaired Student’s t-test.

The G-DRD2 is well distributed at the plasma membrane in HEK 293A cells and showed a rapid increase of green intensity upon the treatment of 10 μM dopamine (Fig. 2D-E, Movie S2). The EC_50_ of the dopamine-induced G-DRD2 response was 27 nM (Fig. 2F), which is much lower than that of R-DRD1 (Fig. 1F). In fact, the binding affinity of dopamine is known to be greater for DRD2 than DRD1 (*40*), suggesting that the EC_50_ values of R-DRD1 and G-DRD2 reflect the differential binding affinities of DRD1 and DRD2. The response of G-DRD2 was specific for the treatment of dopamine or quinpirole, but not to the DRD1 specific agonist SKF (Fig. 2G). In addition, the dopamine-induced G-DRD2 response was prevented by a DRD2 specific inhibitor haloperidol (Halo) (Fig. 2G), confirming the specificity of G-DRD2 to report the activation of DRD2.

Recently, the improved green dopamine sensor called GRAB_DA2m_ was reported (*35*) by the group who developed GRAB_DA1m_ (*33*). GRAB_DA2m_ shows 2 to 3-fold higher dynamic range than GRAB_DA1m_ as a result of modifications on the linker sequences and cpFP in the sensor. The dopamine binding affinity of GRAB_DA2m_ was further tuned by mutations on the sequence of DRD2. Thus, GRAB_DA2m_ is a very sensitive dopamine sensor with stronger dopamine binding affinity. However, we think G-DRD2 can report more physiological aspects of DRD2 keeping its natural dopamine binding affinity, and is thus appropriate for applying to the investigation of the DRD1-DRD2 heterodimer.

### Simultaneous and selective monitoring of DRD1 and DRD2 activity in cells, primary neurons, and brain slices

Our multicolor sensors for dopamine receptors, R-DRD1 and G-DRD2, can be applied for simultaneous and selective monitoring of DRD1 and DRD2 in live cells. We prepared HEK 293A cells expressing either R-DRD1 or G-DRD2, then cultured them together from one day before dual-color imaging. In response to quinpirole, we successfully monitored the selective increase of fluorescence intensity in the cell expressing G-DRD2 but not R-DRD1, and the subsequent treatment of SKF increased the red fluorescence of the R-DRD1-expressing cell (Fig. 3A-B, Movie S3). On these dishes, the cells expressing either R-DRD1 or G-DRD2 showed an increased red or green intensity in response to 10 μM of dopamine, which was completely prevented by pretreatment of SCH or Halo respectively (Fig. 3C-F). The dopamine-induced fluorescent intensity of R-DRD1 or G-DRD2 was also selectively decreased by the subsequent treatment of each inhibitor (Fig. 3C-F). Finally, in the cells expressing both R-DRD1 and G-DRD2, the dopamine-induced responses were successfully monitored without spectral overlapping (fig. S3). These data suggest that the R-DRD1 and G-DRD2 can simultaneously and selectively report the real-time activity of DRD1 and DRD2 in live cells.

**Figure 3.**
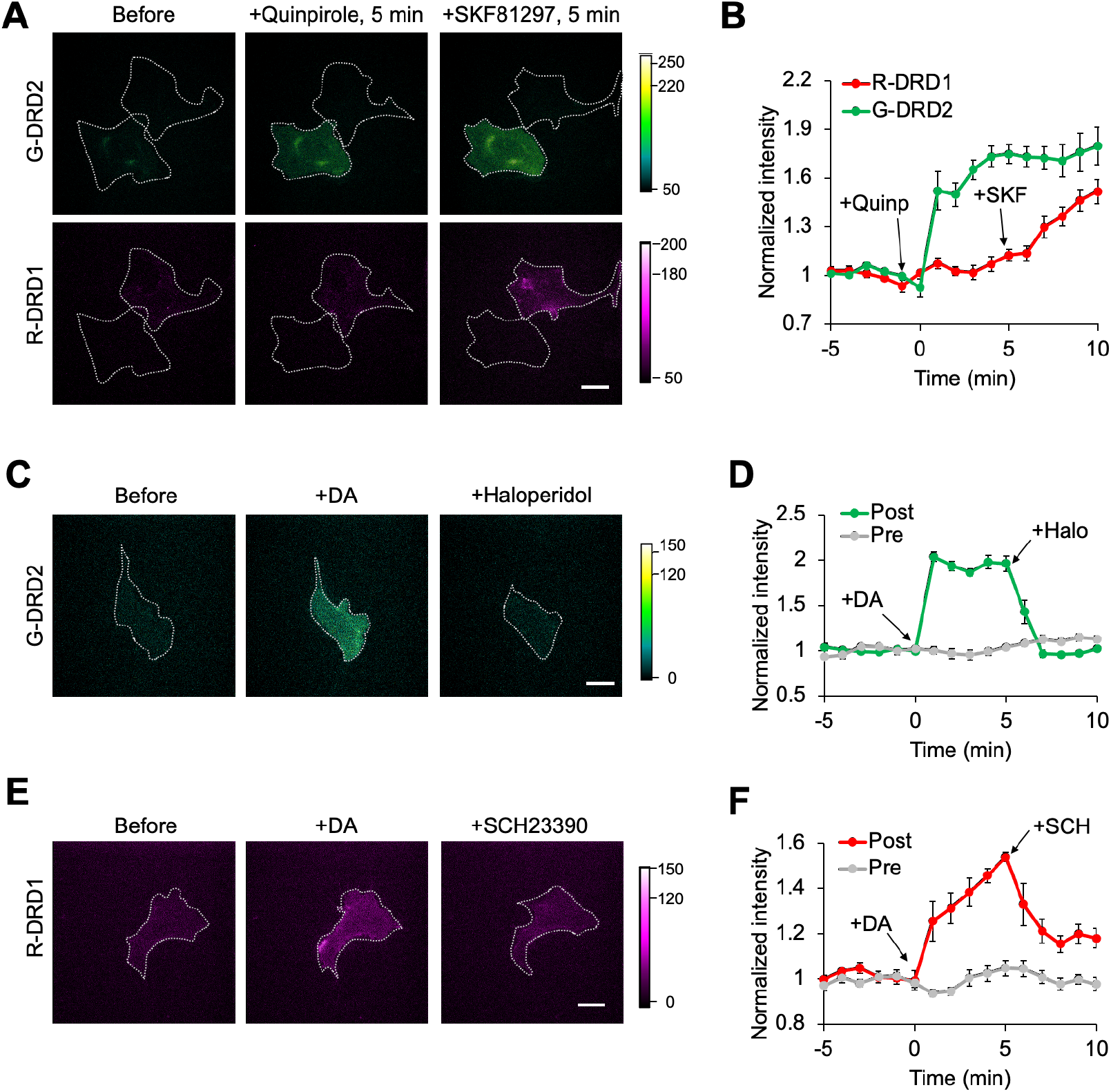
Specific monitoring of DRD1 or DRD2 activity by R-DRD1 or G-DRD2. (**A-B**) Representative dual color images (**A**) and time courses (**B**) of R-DRD1 and G-DRD2 in the same culture dish upon the sequential treatment of quinpirole (10 μM, n = 8) and SKF81297 (10 μM, n = 7). Data are shown as means ± s.e.m. Scale bar, 20 μm. (**C-D**) Representative images (**C**) and time course response (**D**) of G-DRD2 upon the sequential treatment of dopamine (10 μM) and haloperidol (10 μM) (Post-treatment of antagonist, n = 9). The graph also presents the time course of G-DRD2 upon the addition of dopamine (10 μM) in the cells pretreated of haloperidol (10 μM) (Pre, n = 7). Data are shown as means ± s.e.m. Scale bar, 20 μm. (**E-F**) Representative images (**E**) and time course response (**F**) of R-DRD1 upon the sequential treatment of dopamine (10 μM) and SCH23390 (10 μM) (Post-treatment of antagonist, n = 6). The graph also shows the time course of R-DRD1 upon the addition of dopamine (10 μM) in the cells pretreated of SCH23390 (10 μM) (Pre, n = 12). Data are shown as means ± s.e.m. Scale bar, Scale bar, 20 μm.

To validate the multicolor dopamine receptor sensors in neurons, we infected adeno-associated viruses (AAVs) containing R-DRD1 or G-DRD2 in primary cortical neurons isolated from rat embryos (E18) (Fig. 4A). We confirmed the membrane localization of both R-DRD1 and G-DRD2 in the primary cortical neurons (fig. S4). Furthermore, we elaborated on the feasibility of measuring the dopaminergic responses in three-dimensional (3D) neural tissues. The primary cortical neurons cultured within collagen gel (*41, 42*) showed significantly increased fluorescent intensity in response to 100 μM dopamine: a 1.8-fold increase for R-DRD1 within 2 min, a 2.3-fold increase for G-DRD2 within 1 min (Fig. 4B-E, Movies S4 and S5). These data indicate that we can monitor the activity of both DRD1 and DRD2 in live neurons cultured in 3D scaffolds.

**Figure 4.**
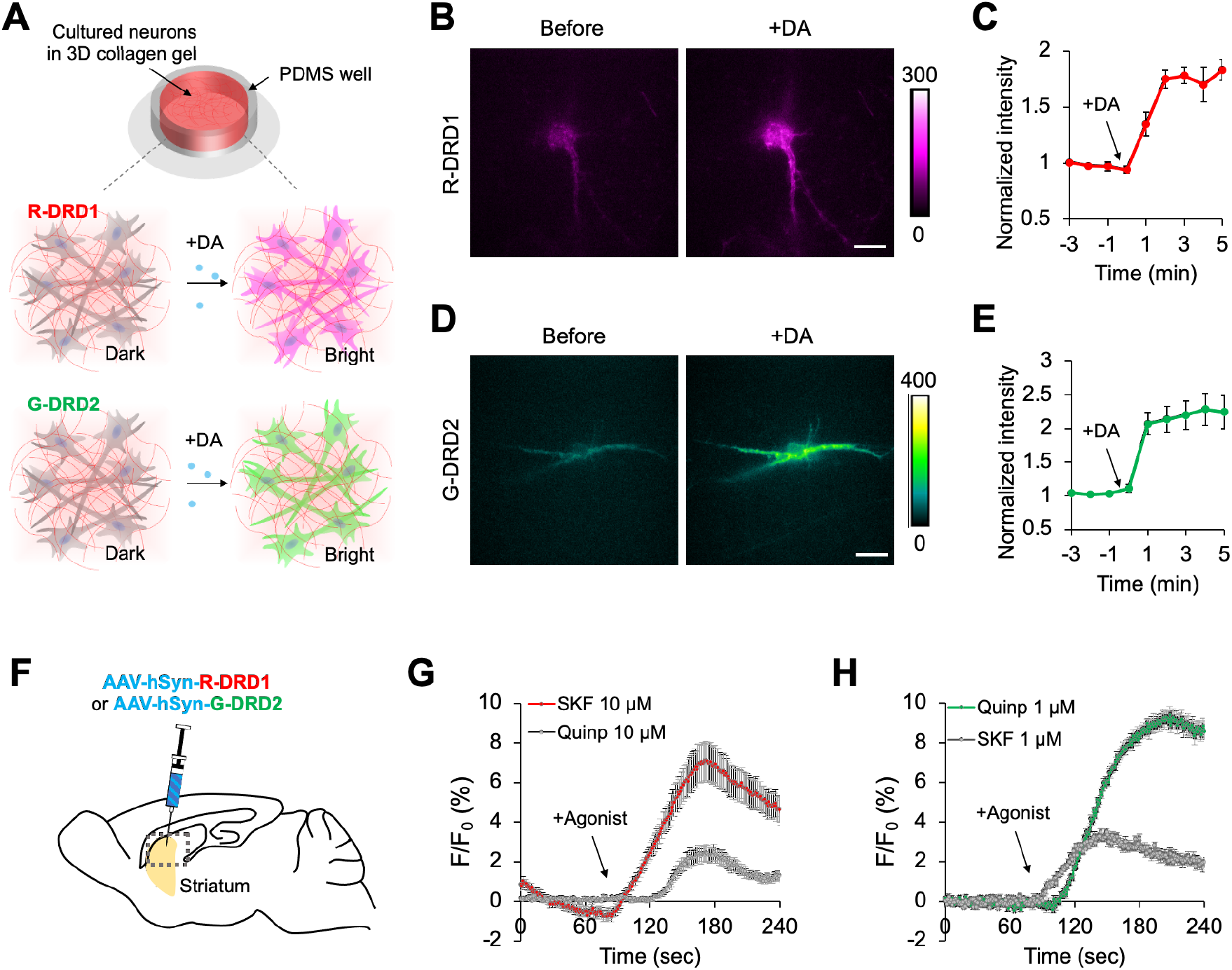
Specific monitoring of DRD1 and DRD2 activity in primary neurons and mouse brain tissues. (**A**) Schematic of rat primary cortical neurons in 3D collagen gel system and the responses of R-DRD1 or G-DRD2 upon the treatment of dopamine. (**B-C**) Representative images (**B**) and time course response of R-DRD1 (**C**) upon the treatment of 100 μM dopamine in primary cortical neurons cultured in 3D collagen gel (n = 5). Data represent the mean value ± s.e.m. (**D-E**) Representative images (**D**) and time course response of G-DRD2 (**E**) upon the treatment of 100 μM dopamine in primary cortical neurons cultured in 3D collagen gel (n = 3). Data represent the mean value ± s.e.m. (**F**) Schematic of mouse brain striatum region injected with viruses containing AAV-hSyn-R-DRD1 or AAV-hSyn-G-DRD2. Normalized fluorescent intensity of R-DRD1 (**G-H**) Specific monitoring of DRD1 or DRD2 activity by R-DRD1 **(G)** or G-DRD2 (**H**) in mouse brain tissues in response to 1 or 10 μM of SKF or Quinp as indicated (n = 14 for **G**, n = 10 for **H**). Data represent the mean value ± s.e.m.

In addition, we investigated whether R-DRD1 and G-DRD2 can selectively visualize the activation of DRD1 and DRD2 in mouse brain slices. We injected the AAVs containing R-DRD1 or G-DRD2 into mouse dorsal striatum and prepared acute brain slices expressing R-DRD1 or G-DRD2. In consistent with the findings from cell experiments, our *ex vivo* imaging of red fluorescence demonstrated that R-DRD1 was preferentially activated by SKF, a DRD1-specific agonist, but significantly less by a DRD2-specific agonist quinpirole (Fig. 4F-G). On the other hand, *ex vivo* imaging of green fluorescence demonstrated that G-DRD2 was much preferentially activated by Quinp, but significantly less by SKF. These findings suggest that R-DRD1 and G-DRD2 can be applied for *ex vivo* brain systems to selectively monitor the activation of DRD1 and DRD2, and more importantly, these sensors help segregate the activity of DRD1 and DRD2.

### Differential receptor crosstalk in the DRD1-DRD2 heterodimer at different dopamine concentrations

Utilizing the R-DRD1 and G-DRD2 sensors, we next investigated whether DRD1 or DRD2 activity in the DRD1-DRD2 heterodimer can be modulated by receptor crosstalk. First, we prepared the cells co-expressing R-DRD1 and DRD2 (Fig. 5A), and confirmed the formation of the heterodimer by co-immunoprecipitation (fig. S5A). The dopamine-induced response of R-DRD1 in the heterodimer was not greatly altered by nanomolar levels of low dopamine concentrations (Fig. 5B). However, the activation of R-DRD1 in the heterodimer was significantly reduced by the treatment of dopamine at concentration of 10 μM or more (Fig. 5B). Interestingly, tonic and phasic dopamine release are suggested in the ranges of nanomolar and micromolar dopamine concentrations (*39, 40, 43, 44*). Thus, our data imply that the DRD1 activity in the DRD1-DRD2 heterodimer may be differentially regulated at tonic and phasic states.

**Figure 5.**
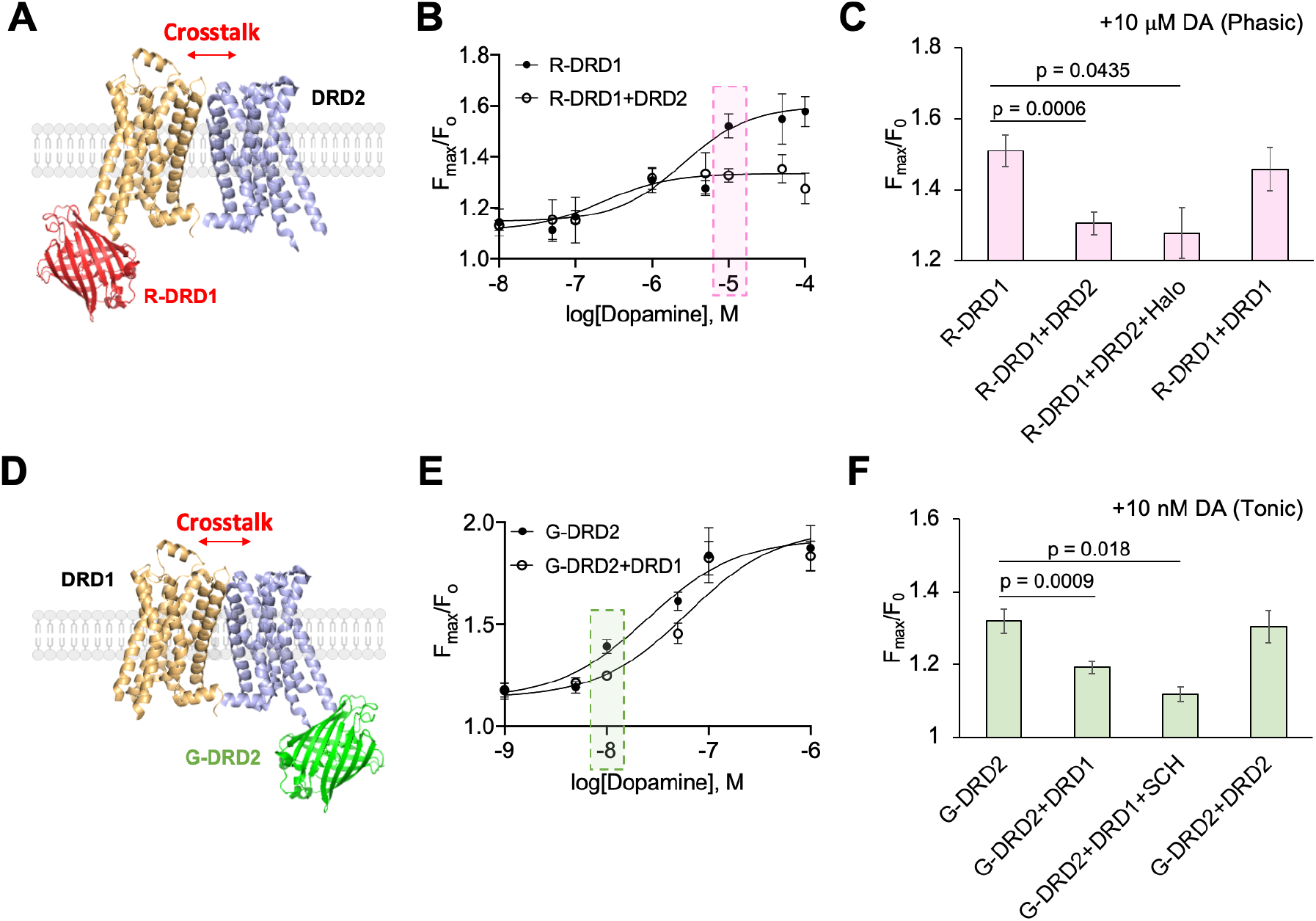
Differential receptor crosstalk in the DRD1-DRD2 heterodimer upon different dopamine concentrations. **(A)** Experimental schematic for monitoring of DRD1 activity in the R-DRD1-DRD2 heterodimer. **(B)** *F*_max_ /*F*_0_ of R-DRD1 or R-DRD1-DRD2 heterodimer in response to different dopamine concentrations. Each dot represents the *F*_max_ /*F*_0_ of R-DRD1-DRD2 at different dopamine concentrations from more than 4 cells. The *F*_max_ /*F*_0_ of R-DRD1-DRD2 is significantly lower than the one of R-DRD1 at the dopamine concentrations of 10 μM (pink box). Data represent the mean value ± s.e.m. **(C)** *F*_max_ /*F*_0_ in response to 10 μM of dopamine treatment, in the cells expressing R-DRD1 (n = 35), R-DRD1 and DRD2 (n = 29), R-DRD1 and DRD2 with the pretreatment of haloperidol (10 μM) (n = 6), R-DRD1 and DRD1 (n = 8). Data represent the mean value ± s.e.m. and p values were calculated by Student’s unpaired t-test. **(D)** Experimental schematic for monitoring of DRD2 activity in the DRD1-G-DRD2 heterodimer. **(E)** *F*_max_ /*F*_0_ of G-DRD2 or DRD1-G-DRD2 in response to different dopamine concentrations. Each dot represents the *F*_max_ /*F*_0_ of DRD1-G-DRD2 at different dopamine concentrations from more than 6 cells. The *F*_max_ /*F*_0_ of DRD1-G-DRD2 is significantly lower than the one of G-DRD2 at the dopamine concentrations of 10 nM (green box). Data represent the mean value ± s.e.m. **(F)** *F*_max_ /*F*_0_ in response to 10 nM of dopamine treatment, of the cells expressing G-DRD2 (n = 43), G-DRD2 and DRD1 (n = 42), G-DRD2 and DRD1 with the pretreatment of SCH23390 (10 μM) (n = 5), G-DRD2 and DRD2 (n = 8). Data represent the mean value ± s.e.m. and p values were calculated by Student’s unpaired t-test.

This selective inhibition of R-DRD1 in the heterodimer after the treatment of 10 μM dopamine, but not at 10 nM concentration (fig. S6A), was not prevented by Halo, a DRD2 specific inhibitor (Fig. 5C), implying that the reduced DRD1 activity within the heterodimer is not due to the activity of DRD2, but possibly because of the physical interaction between DRD1 and DRD2. To exclude the possibility that the DRD1 inhibition is simply due to the increased density of receptors, we introduced R-DRD1 and DRD1 into same cell, but no significant alteration in the R-DRD1 response was detected (Fig. 5C). These results suggest that the DRD1 activity is inhibited within the DRD1-DRD2 heterodimer, particularly in response to phasic state of micromolar dopamine concentrations.

We next prepared the cells co-expressing G-DRD2 and DRD1 (Fig. 5D), and confirmed the generation of the heterodimer (fig. S5B). We then applied different concentrations of dopamine to investigate the receptor crosstalk effect on DRD2 activation within the DRD1-DRD2 heterodimer. Interestingly, the G-DRD2 response in the heterodimer was selectively inhibited at low dopamine concentration (10 nM), but not at micromolar high dopamine concentrations (Fig. 5E, fig S6B). This inhibition of G-DRD2 in the heterodimer at 10 nM dopamine was not prevented by a DRD1 inhibitor SCH nor co-expression of DRD2 (Fig. 5F). Taken together, the DRD1-DRD2 heterodimer modulates the dopamine-induced activation of DRD1 and DRD2 at different dopamine concentrations: DRD1 activation is inhibited at micromolar phasic concentrations (10 μM or more) whereas DRD2 activation is reduced at nanomolar tonic concentration (10 nM).

### The DRD1-DRD2 heterodimer differentially modulates the cAMP levels at different dopamine concentrations

After binding to dopamine, DRD1 interacts with Gs protein and activates adenylyl cyclase for producing cAMP, while DRD2 activates Gi activity which inhibits AC lowering the cAMP levels (*4*). It has been suggested that the DRD1-DRD2 heterodimer can activate another G alpha subunit, Gq (*18, 45*), which further induces calcium-related signaling (*16, 21, 46*). We also confirmed, utilizing a fluorescent calcium sensor jRGECO1a (*36*), that the DRD1-DRD2 heterodimer can increase intracellular calcium levels in a dopamine dose-dependent manner (fig. S7). However, other study suggested that the calcium-related signaling pathways by the DRD1-DRD2 heterodimer may be mediated by multiple synergistic effects (*47*), suggesting that the downstream signaling mechanisms of DRD1-DRD2 heterodimer are complex and still require more investigations.

While calcium-related downstream signaling of the DRD1-DRD2 heterodimer has been studied (*16, 18, 45*), it was unclear how their original functions on cAMP signaling via Gs or Gi are regulated in the heterodimer. In particular, we discovered differential receptor crosstalk from the DRD1-DRD2 heterodimer (Fig. 5). Thus, utilizing a cAMP sensor GloSensor (*38*), we further explored whether and how the heterodimer modulates the cAMP levels upon different dopamine concentrations. We first confirmed that activated DRD1 increases the level of cAMP (Fig. 6A, yellow bars), whereas DRD2 activation induced the decreased cAMP level (Fig. 6B). Upon the treatment of 10 nM dopamine, the DRD1-DRD2 heterodimer induces similar level of cAMP to DRD1 (Fig. 6A), possibly due to the selective inhibition of DRD2 activity in the heterodimer at this nanomolar dopamine level (Fig. 6C and Fig. 5E). In contrast, at 10 μM dopamine, the cAMP level by DRD1-DRD2 heterodimer was significantly lower than the one by DRD1 (Fig. 6A), suggesting the selective inhibition of DRD1 at micromolar dopamine concentration (Fig. 6D and Fig. 5B). These results are consistent with the receptor crosstalk data in the DRD1-DRD2 heterodimer (Fig. 5). Therefore, differential receptor crosstalk in the DRD1-DRD2 heterodimer depending on dopamine concentrations can further modulate their original downstream cAMP signaling via Gs or Gi proteins.

**Figure 6.**
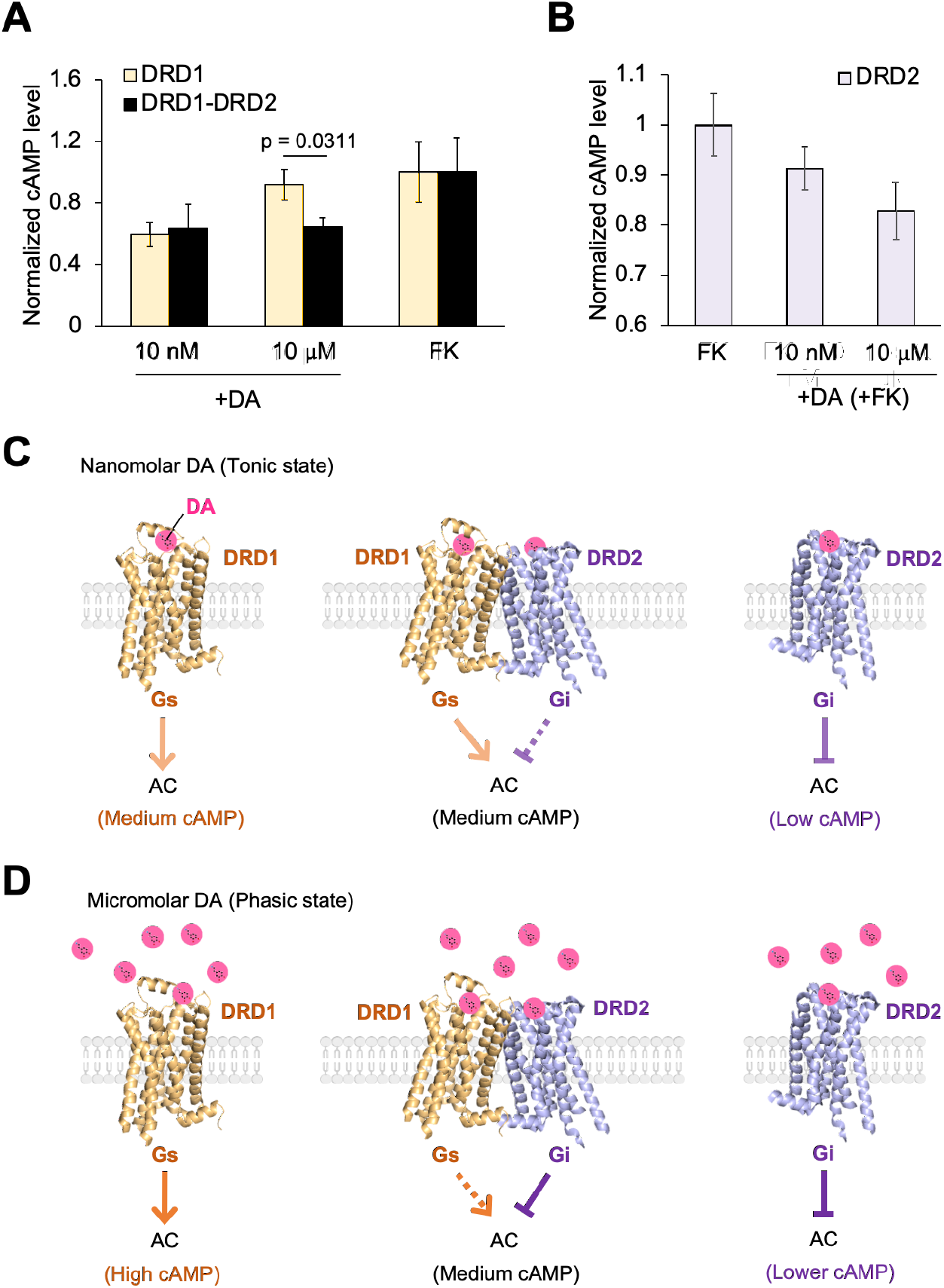
The DRD1-DRD2 heterodimer differentially modulates the cAMP levels at different dopamine concentrations. (**A**) Normalized cAMP levels, detected by GloSensor, in HEK293A cells expressing DRD1 (yellow bars) or DRD1-DRD2 (black bars) in response to dopamine (10 nM or 10 μM) or forskolin (10 μM) as a positive control (n = 4). Data represent the mean value ± s.e.m. and p values were calculated by Student’s unpaired t-test. (**B**) Normalized cAMP levels in the HEK293A cells expressing DRD2 upon the incubation of forskolin (10 μM) without or with dopamine (10 nM or 10 μM) (n = 4, 6, 7). Data represent the mean value ± s.e.m. (**C**) Model for tonic state. At nanomolar dopamine concentration, the DRD2-induced Gi function lowering cAMP level is selectively inhibited in the DRD1-DRD2 heterodimer. (**D**) Model for phasic state. At micromolar dopamine concentration, the DRD1-induced Gs function, i.e. cAMP production, is selectively reduced in the DRD1-DRD2 heterodimer. This results in the significantly lower cAMP level in the cells expressing the heterodimer comparing to the one in DRD1-expressing cells.

## Discussion

In this study, we developed multicolor dopamine receptor sensors, R-DRD1 and G-DRD2, for the simultaneous and selective monitoring of DRD1 and DRD2 activities in live cells, primary neurons, and mouse brain tissues (Fig. 1–4). In particular, R-DRD1 and G-DRD2 sensors can specifically detect the activation status of DRD1 and DRD2 with their natural property, as the EC_50_ of R-DRD1 and G-DRD2 are 2.2 μM and 27 nM, respectively, similar to their natural dopamine binding affinity (*44, 48, 49*). Importantly, these EC_50_ values are in the physiological ranges of phasic and tonic dopamine release. Thus, the multicolor DRD1 and DRD2 sensors are novel tools for the investigation of physiological roles of the DRD1, DRD2, and furthermore DRD1-DRD2 heterodimer upon phasic and tonic dopamine signals.

Balance between tonic and phasic dopamine release is important for background homeostasis as well as initiation of dopamine signaling pathways (*50, 51*). In response to nanomolar ranges of tonic dopamine signals, activated DRD2 is known to inhibit the cAMP signaling pathways and mediate the function of inhibitory GIRK channels (*2*). As DRD2 is also distributed at presynaptic axons, and this inhibitory DRD2 autoreceptor can negatively regulate the firing patterns of dopaminergic neurons (*52*). In contrast, micromolar phasic dopamine bursts are suggested to activate DRD1 more efficiently than DRD2 (*53*), allowing local and transient activation of DRD1 and thus efficient transduction of the positive dopamine signaling pathway.

It has been difficult to directly monitor the status of receptor activation, thus the activation patterns of DRD1, DRD2, and DRD1-DRD2 heterodimer at tonic and phasic dopamine signals can be only suspected by measuring their downstream signaling pathways (*16, 54–56*). As we developed multicolor dopamine receptor sensors, R-DRD1 and G-DRD2, which can specifically report the activation status of DRD1 and DRD2, the functional crosstalk in the DRD1-DRD2 heterodimer can be directly measured for the first time (Fig. 5). In particular, we wondered how their activities are regulated by the receptor crosstalk in response to the physiological ranges of phasic and tonic dopamine signals. Surprisingly, we discovered differential receptor crosstalk within the DRD1-DRD2 heterodimer upon different dopamine concentrations: the DRD1 activity in the heterodimer is inhibited only at micromolar (phasic) dopamine concentrations, whereas the DRD2 activity is selectively inhibited at nanomolar (tonic) levels of dopamine.

We further confirmed that the differential receptor crosstalk in the DRD1-DRD2 heterodimer at tonic and phasic dopamine levels modulates their original downstream pathways (Fig. 6). At nanomolar tonic dopamine concentration, DRD2 can induce the reduction of cAMP levels, however the DRD2-mediated Gi function is selectively inhibited in the DRD1-DRD2 heterodimer, thus the cAMP level by the heterodimer is similar to the one by DRD1 (Fig. 6C). In contrast, at micromolar phasic dopamine concentration, DRD1 can induce high cAMP level via Gs function, but the DRD1 activity is selectively inhibited in the DRD1-DRD2 heterodimer, hence the cAMP level by the heterodimer is significantly lower than the one by DRD1 (Fig. 6D). Consequently, the cAMP level changes in response to phasic and tonic dopamine signals are much attenuated in the cells expressing the DRD1-DRD2 heterodimer, which may further influence downstream signaling pathways, such as PKA, DARPP-32, MAPK, and specific gene expressions (*47, 55*).

Indeed, the alteration of the DRD1-DRD2 population can be observed both in physiological and pathological conditions. For example, the DRD1-DRD2 heterodimer was increased in striatum from older mice (*45*), which may explain the vulnerability of dopamine-related disorders at aged conditions. In addition, a higher density of DRD1-DRD2 heterodimer population was observed in the macaque brain in the Parkinson’s disease model (*57*). These evidences imply that the DRD1-DRD2 heterodimer is important in dopamine-related diseases and the dopamine functions at aged conditions (*58*).

Recent studies proposed that drug addiction can be also related to the DRD1-DRD2 heterodimer (*14, 56, 59, 60*). Comparing to the control group, cocaine addicted patients showed a significant increased population of the DRD1-DRD2 heterodimer in the striatum (*60*). This alteration in dopamine receptor populations may be due to the sustained high concentration of dopamine in the synaptic cleft by the cocaine-mediated blockade of the dopamine transporter (*61*). The sustained high dopamine levels can prolong the activation status of dopamine receptors and generate excessive downstream signaling (*62, 63*), which may cause an increased population of DRD1-DRD2 heterodimer to attenuate the response induced by these strong dopamine signals. Transient formation of the DRD1-DRD2 heterodimer may downregulate abnormal dopamine transmission, however chronic enhancement of the DRD1-DRD2 heterodimer may diminish the dopamine-mediated signaling at postsynapses and eventually disturb normal signal transmission in dopaminergic brain circuits. Thus, addictive cocaine administration can dysregulate the normal dopamine functions.

In summary, we developed novel DRD1 and DRD2 sensors, R-DRD1 and G-DRD2, and discovered differential receptor crosstalk in the DRD1-DRD2 heterodimer upon different dopamine concentrations in the ranges of phasic and tonic dopamine signals. The receptor crosstalk in the heterodimer further modulates their downstream cAMP levels, which consequently attenuates the changes of cAMP levels by DRD1-DRD2 heterodimer. Our results propose a novel perspective on the roles of the DRD1-DRD2 heterodimer for aging, psychological disorders and drug addiction. These will be further investigated with R-DRD1 and G-DRD2 in 3D neural culture or brain organoid, as we elaborated on the feasibility of measuring the DRD1 and DRD2 responses in 3D neural cultures and brain tissues. Recently, in vivo imaging of dLight1.1 in DRD1- and DRD2-specific cre mice revealed that cell-type-specific modulation of PKA by dopamine is crucial for learning (*64*). We believe simultaneous in vivo imaging of DRD1 and DRD2 can provide further critical information of the functions of DRD1, DRD2, and DRD1-DRD2 heterodimer. Therefore, our multicolor R-DRD1 and G-DRD2 sensors will be applied for in vivo multicolor imaging to real their distinct roles in physiological and pathophysiological conditions. Direct observation of receptor activity using conformational change based multicolor GPCR sensors in live cells provides uncovered information about crosstalk effect during heterodimer complex, and this approach will be further applied to investigate the function of other GPCR heterodimers.

## Acknowledgments

This work is supported by KIST Institutional Grant (2E30963), Brain Research Program through the National Research Foundation of Korea (2017M3C7A1043842), Samsung Research Funding & Incubation Center of Samsung Electronics (SRFC-TC2003-02) (J.S.).

## Author Contributions

J.S., H.K, M.H.N, N.C designed research; H.K., M.H.N., S.J., H.L. performed experiments; J.S., H.K., M.H.N, S.J., H.L., S.J.O., J.K., N.C. analyzed data; J.S., H.K., M.H.N., N.C. wrote the manuscript.

## Materials and Methods

### Plasmids

Human DRD1 and DRD2 in pcDNA vectors were achieved using the PRESTO-Tango kit (Addgene). To generate R-DRD1, the cpmApple module from jRGECO1a was amplified by PCR and inserted by In-Fusion (Clontech) to the ICL3 region of human DRD1 in the pcDNA5 vector. For optimizing the linker sequences in the insertion sites, site saturation mutagenesis was performed by PCR with NNK codon sequence. The G-DRD2 sensor was generated by inserting the cpGFP module from dLight1.1 to the ICL3 of human DRD2 in the pcDNA5 vector. pGlo-22F was achieved from Promega for measuring the cAMP levels in live cells. For the expression in primary neurons and brain tissues, R-DRD1 and G-DRD2 were subcloned into a pAAV-hSyn vector, and the adenoviruses were produced by KIST virus facility. pCMV-dLight1.1, pcDNA-GRAB-DA1m and pGP-CMV-NES-jRGECO1a were obtained from Addgene.

### Cell Culture, transfection, and reagents

The human embryonic kidney 293A (HEK293A) cell line was maintained in DMEM supplemented with 10% fetal bovine serum (Hyclone), 1 unit per ml penicillin, 100 μg per ml streptomycin and 100 μM MEM Non Essential Amino Acids Solution (Gibco). Cell culture reagents were purchased from Hyclone. Cells were cultured in a humidified 95% air, 5% CO_2_ incubator at 37C. Lipofectamine™ 2000 reagent (Invitrogen) was used for the transfection according to the manufacturer’s protocol. Dopamine hydrochloride, quinpirole, R(+)-SCH-23390 hydrochloride and haloperidol were purchased from Sigma-Aldrich. SKF81297 and forskolin were purchased from Tocris.

### Isolation and 3D culture of rat primary cortical neurons

Pregnant Sprague Dawley rats (E18) were purchased from SAMTAKO and sacrificed for primary cortical neurons, following the previous protocol (*41, 42*). All procedures were conducted according to the animal welfare guidelines approved by the Institutional Animal Care and Use Committee of the Korea Institute of Science and Technology. Briefly, the entire cortex was dissected out as an intact structure from the brains of embryos under the dissection microscope. The cortex was treated with papain (Miltenyi Biotec) and serially triturated.

Dissociated cells were counted and seeded at a density of 4 x 10^6^ cells ml^-1^ in the neutralized collagen solution adjusted at 0.25% [w/v] of final concentration. Cells were cultured in medium consisting of neurobasal media supplemented with 2% [v/v] B27-supplement (Invitrogen), 2 mM Glutamax-I (Gibco) and 1% [v/v] penicillin-streptomycin (Gibco) at 37 °C in a 5% CO_2_ humidified incubator. One-half of the medium was replaced with a fresh culture medium every 2-3 days.

### Live-cell image acquisition

Live-cell imaging was performed in a humidified 95% air, 5% CO_2_ and 37°C temperature-controlled chamber (Live Cell Instrument). HEK293A cells expressing dopamine receptor sensors or other constructs were prepared on cover-glass-bottom dishes coated with 10 μg/ml of fibronectin (Gibco). Various concentration of dopamine or other ligands were added to the cells. Cells were monitored using the 100x objective lens for indicated times. Images were collected by a Nikon Ti-E inverted microscope and a cooled charge-coupled device camera using NIS software (Nikon). G-DRD2 images were collected using a 482DF35 excitation filter, a 506DRLP dichroic mirror, and 536DF40 emission filter. R-DRD1 images were collected using a 562DF40 excitation filter, a 593DRLP, a 641DF75 emission filter. A neutral-density filter was used to control the intensity of the excitation light. For the response measurement, the images of G-DRD2 (ex: 482DF35, em: 536DF40) and R-DRD1 (ex: 562DF40, em: 641DF75) at each time point were achieved with exposure time of 50 ms with ND 8 filter. The fluorescence intensity of non-transfected cells was quantified as the background signal and subtracted from the fluorescence signals from transfected cells. The pixel-by-pixel fluorescent intensity images were calculated based on the background-subtracted fluorescence intensity images of G-DRD2 and R-DRD1 by the NIS program to allow the quantification and statistical analysis.

### Ex vivo imaging of mouse brain slices

AAV-hSyn-R-DRD1 or AAV-hSyn-G-DRD2 viruses were injected into the dorsal striatum of 6 to 8-week old C57BL/6 mice (ML: +/-2 mm, AP: +0.7 mm, DV: −3 mm from bregma). After 2 to 3 weeks, animals were deeply anesthetized with isoflurane, followed by decapitation. The brains were removed from the skull and placed in an ice-cold oxygenated (95% O_2_ and 5% CO_2_) artificial cerebrospinal fluid (ACSF; 130 NaCl, 24 NaHCO_3_, 3.5 KCl, 1.25 NaH_2_PO_4_, 1 CaCl_2_, 3 MgCl_2_ and 10 glucose (in mM), pH 7.4). 300-μm thick coronal slices of striatal tissue was prepared with vibrating microtome (DSK model PRO7) and incubated in recording solution (ACSF; 130 NaCl, 24 NaHCO_3_, 3.5 KCl, 1.25 NaH_2_PO_4_, 1.5 CaCl_2_, 1.5 MgCl_2_ and 10 glucose (in mM), pH 7.4) at room temperature for at least 1 hour before recording. Region of interests (ROIs) were defined under 20x water-immersion objective lens with fluorescent filters. Each ROI was imaged using Image workbench (Indec biosystems) with a scanning speed of 1 frame per second. SKF81297 (1, 10 μM) and Quinpirole (1, 10 μM) were applied for 150 sec in the bath after at least 5 min for allowing the stable baseline. After recording with the drugs, KCl (20 μM) was treated to induce striatal axon depolarization for the confirmation of dopamine receptor sensors. Prior to quantification of the change in the fluorescence intensity, the baseline (before drug treatment and 100 sec after the termination of drug application) was appropriately adjusted, then the (F-F_0_)/F_0_ (%) was calculated from each trace.

### GloSensor cAMP assay

For the GloSensor cAMP assay, pGlo-22F and DRD plasmids (DRD1, DRD2, or DRD1 and DRD2) were co-transfected in HEK293A cells 24-36 hours. The transfected cells were harvested and seeded in 96-well plates at a density of 2 × 10^4^ cells for 24 hours. On the day of cAMP assay, the media is removed and D-luciferin sodium salt (GoldBio, 150 μg/ml) in 20 mM HEPES containing HBSS buffer (pH 7.4) was added to the wells and incubated for 30 min at 37 °C. After the treatment of dopamine (10 nM or 10 μM) or forskolin (10 μM), luminescence signal was measured using Synergy H1 (Bio Tek) at room temperature.

### Statistical analysis

To determine the p values between experimental and control groups, statistical analysis was performed using unpaired Student’s t-test at GraphPad Prism 8 or Excel, and the values are shown as mean ± standard error of the mean (s.e.m.).

